# Pathogenicity and intestinal barrier disruptive ability of *Malassezia furfur* in an alternative model host *Caenorhabditis elegans* is partially alleviated by *Lacticaseibacillus rhamnosus*

**DOI:** 10.1101/2024.10.17.618914

**Authors:** Chiho Kishida, Ayano Tsuru, Satoka Takabayashi, Xueyang Wu, Yoshihiko Tanimoto, Eriko Kage-Nakadai

**Affiliations:** Graduate School of Human Life and Ecology, Osaka Metropolitan University, Osaka, Japan; Graduate School of Human Life Science, Osaka City University, Osaka, Japan; Department of Food and Nutrition, Faculty of Human Life Science, Osaka City University, Osaka, Japan; Institute for Life and Medical Sciences, Kyoto University, Kyoto, Japan

## Abstract

*Malassezia furfur* is associated with various diseases; however, the mechanisms underlying its pathogenicity remain largely unknown. In the present study, *Caenorhabditis elegans* was used as the model host to evaluate *M. furfur* pathogenicity. Additionally, effects of lactic acid bacteria against *M. furfur* pathogenicity were evaluated. Compared to *Escherichia coli* OP50 (OP, control), both live and heat-killed *M. furfur* reduced the lifespan and body size of *C. elegans*, although heat-killed *M. furfur* was less effective than live *M. furfur* in lifespan shortening. Furthermore, unlike heat-killed *M. furfur,* live *M. furfur* disrupted the nematode intestinal barrier. *nsy-1* and *sek-1* loss-of-function mutants were susceptible to *M. furfur*, suggesting their involvement in the defense against *M. furfur* infection. Expression of genes involved in host defense and of those coding for C-type lectin domain-containing proteins and antimicrobial peptides was upregulated in *M. furfur*-infected *C. elegans*. *Lacticaseibacillus rhamnosus* (LR) significantly ameliorated lifespan shortening and body size reduction in *M. furfur*-infected *C. elegans* and protected against intestinal barrier disruption, suggesting that LR protects nematodes from *M. furfur* virulence. This study highlights *M. furfur* pathogenicity and intestinal barrier disruptive ability in *C. elegans* and suggests that the *M. furfur* virulence is partially attenuated by LR.

**Importance:** Infection with *Malassezia furfur* shortens the lifespan and disrupts the intestinal barrier in the model host *C. elegans*. The probiotic *Lacticaseibacillus rhamnosus* (LR) attenuates *M. furfur* virulence, thus partially protecting the intestinal tract. Signaling of the innate immune response to *M. furfur* in *C. elegans* is mediated by *nsy-1* and *sek-1*, suggesting that the expression of genes involved in the biological defense response may be regulated downstream of *nsy-1* and *sek-1*. This study enhances our understanding of the diseases associated with *M. furfur* and offers insights into potential preventive and therapeutic methods using probiotics.

## Introduction

*Malassezia* is a dimorphic fungus found in the human skin flora and dominates more than half of all the fungal species on the skin (1). *Malassezia* requires lipids for growth and is predominantly found in lipid-rich areas of the human body, such as the scalp and trunk (2). The fungus is present on the skin under normal conditions and is associated with various cutaneous and systemic diseases, including pityriasis versicolor, folliculitis, seborrheic dermatitis, as well as with sepsis related to deep venous catheters in infants (3–6). Although the exact mechanisms underlying these diseases are not fully understood, hyphal formation in patients with pityriasis versicolor and upregulation of lipases and metabolic product (such as indole) in patients with seborrheic dermatitis has been reported (7–9). Furthermore, *Malassezia* has been found not only on the skin but also in the gut of patients with inflammatory bowel disease, and mucosal lavage samples from patients with Crohn’s disease had increased levels of *Malassezia* compared to healthy controls (10). Nevertheless, interactions between *Malassezia* and its host remain ambiguous and studies to determine its potential as a contaminant are warranted owing to *Malassezia* being a cutaneous microflora (11).

Currently, the genus *Malassezia* comprises 18 species (12). Among these, *Malassezia furfur* has been implicated in various skin diseases, including pityriasis versicolor and seborrheic dermatitis (13). The genus is also the most frequently identified in bloodstream infections, which occur via catheters in immunocompromised patients and neonates (14). Several indoles induced by *M. furfur*, including malassezin, activate the aryl hydrocarbon receptor, a transcription factor that regulates various physiological functions of the host, including the immune system. This receptor is also involved in *Malassezia*-related skin disease (9, 15). *M. furfur* is also present in healthy individuals (16, 17); therefore, identifying the developmental factors of *Malassezia*-related diseases has become more complex.

*Caenorhabditis elegans* is used as an alternative model host for pathogen infection studies owing to its short life cycle and conserved innate immune pathways, such as the mitogen-activated protein kinase (MAPK) and insulin signaling pathways, as well as a wide range of available genetic tools (18, 19). Pathogenic bacteria and fungi, such as *Salmonella typhimurium* and *Candida albicans*, respectively, shorten the lifespan of *C. elegans* (20, 21). In contrast, lactic acid bacteria and *Bifidobacterium* exert probiotic effects that prolong the lifespan of *Salmonella*-infected *C. elegans* (22). Thus, *C. elegans* is a suitable host model for evaluating the protective effects of probiotics against pathogens.

In this study, we aimed to use *C. elegans* as a model host to evaluate the effect of *Malassezia* on the gut and to investigate the mechanisms underlying the innate immune response to it. Additionally, we evaluated the effects of lactic acid bacteria on the host’s protection against *M. furfur* pathogenicity.

## Results

### Pathogenicity of *M. furfur* in wild-type (N2) *C. elegans*

To examine the influence of *M. furfur* infection on the lifespan of *C. elegans*, N2 worms were fed *M. furfur* or *Escherichia coli* OP50 (OP, control). The results showed that N2 worms fed *M. furfur* had significantly shorter lifespans than those fed OP (**Fig. 1A**). To determine whether live *M. furfur* was pathogenic, lifespan assays were conducted with N2 worms fed heat-killed (HK)-OP or HK-*M. furfur*. The survival rate of nematodes in the HK-*M. furfur* group was significantly lower than that of those in the HK-OP group, whereas the survival rate of worms fed HK-*M. furfur* was higher than that of those fed live *M. furfur* (**Fig. 1B**). To characterize the pathogenicity of *M. furfur*, we examined the changes in body size and intestinal barrier damage in worms fed *M. furfur*. Worm projection areas were measured beginning at 4 d of age (the day after the start of feeding *M. furfur*) to 6 d of age as an index of body size. Both *M. furfur*- and HK-*M. furfur*-fed groups showed a significant decrease in body area compared to the OP-fed control group (**Fig. 1C**). Intestinal barrier damage was evaluated using the Smurf assay, in which worms were fed a blue dye and examined for leakage of dye from the intestinal lumen into the body cavity (23). For 6-d-old worms, the percentage of worms with body cavity leakage in the *M. furfur*-fed group was significantly higher than that of those in the other groups. In contrast, no significant differences in intestinal barrier damage were observed among the OP, HK-OP, and HK-*M. furfur* groups (**Fig. 1D and E**).

**Fig 1.**
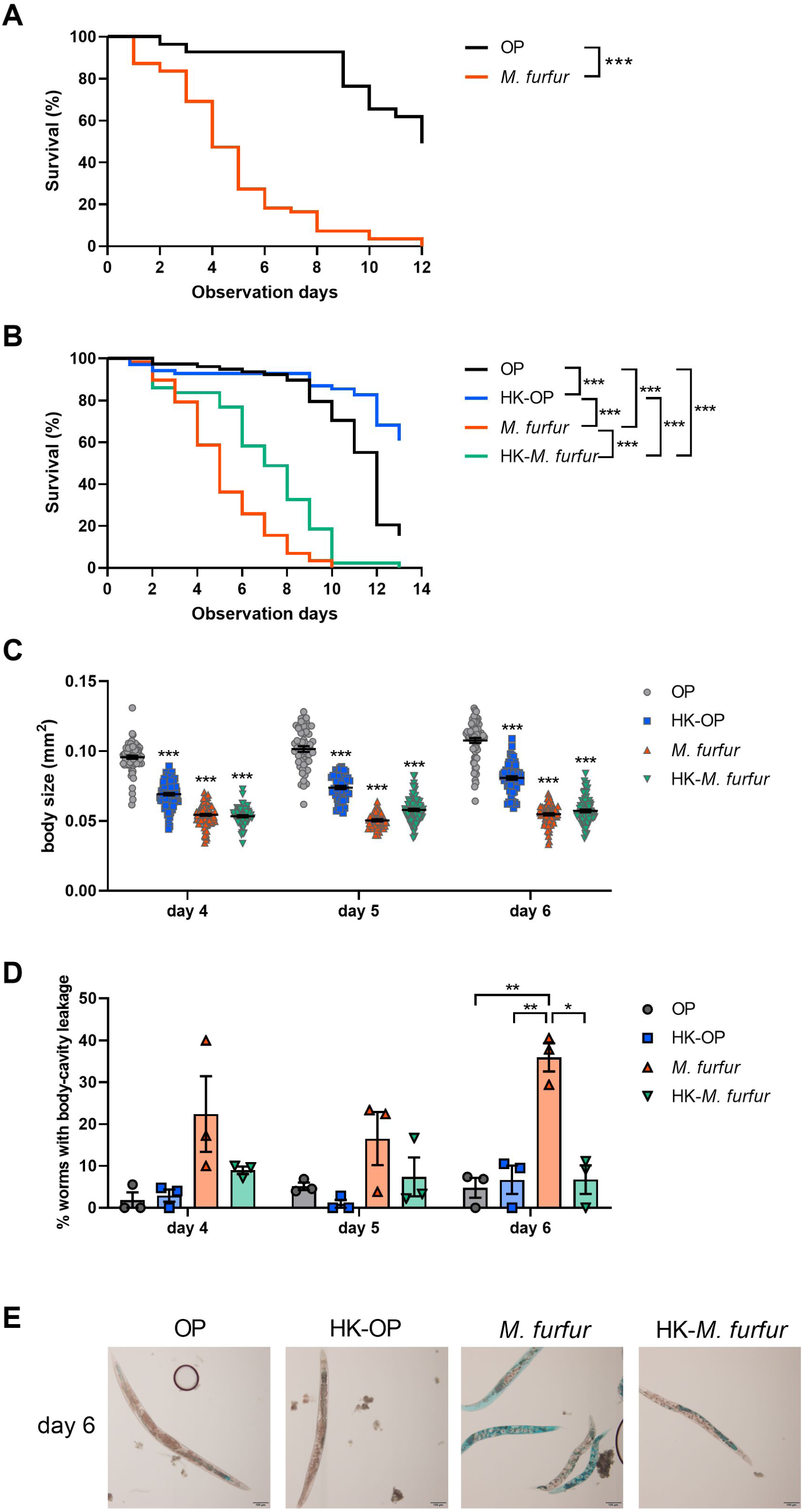
Evaluation of *Malassezia furfur* pathogenicity in *Caenorhabditis elegans*. (A) Survival curves of N2 (wild-type) worms fed *Escherichia coli* OP50 (OP, n = 55) or *M. furfur* (*M. furfur*, n = 55). (B) Survival curves of N2 worms fed live OP or live *M. furfur*, and heat-killed (HK)-OP or HK-*M. furfur* (OP, n = 78; HK-OP, n = 69; *M. furfur*, n = 58; HK-*M. furfur*, n = 43). For (A) and (B), young adult worms (3-d-old) represent day 0 of observation. Survival rates were calculated using the Kaplan–Meier method and asterisks indicate a significant difference compared to OP-fed worms (control) using the log-rank (Mantel-Cox) test. ***, P < 0.001. (C) Body size of worms fed OP, HK-OP, *M. furfur*, or HK-*M. furfur* from 4 to 6 d of age. Asterisks indicate significant differences compared to OP-fed worms (control). (D) The intestinal barrier function was assessed by examining the percentage of worms showing leakage of dye into the body-cavity (Smurf assay). Worms fed OP, HK-OP, *M. furfur*, or HK-*M. furfur* were examined from 4 to 6 d of age. For (C) and (D), results are shown as individual plots and means ± standard error of mean (SEM). Statistical analysis was performed with one-way ANOVA and Tukey’s multiple comparison tests. ***, P < 0.001; **, P < 0.01; *, P < 0.05. (E) Representative images of 6-d-old worms fed OP, HK-OP, *M. furfur*, or HK-*M. furfur* and stained with blue dye. Scale bar, 100 µm.

### Effect of *M. furfur* infection on MAPK mutants

To investigate whether the innate immune response against *M. furfur* in *C. elegans* is mediated by MAPK signal transduction pathways, lifespan assays were performed with *M. furfur*-fed loss-of-function mutants of the *nsy-1*, *sek-1*, *pmk-1*, *mkk-4*, and *jnk-1* genes that encode kinases in the p38 or c-Jun N-Terminal Kinase (JNK) MAPK cascade (24, 25). The survival rates of *nsy-1* and *sek-1* loss-of-function mutants were significantly lower than those of the N2 (control) worms (**Fig. 2A and B, S1 and 2**). In contrast, the survival rates of *pmk-1* and *jnk-1* loss-of-function mutants did not differ significantly from the controls, while the survival rate of the *mkk-4* loss-of-function mutant was higher than that of the controls. (**Fig. 2B and C, S2**).

**Fig 2.**
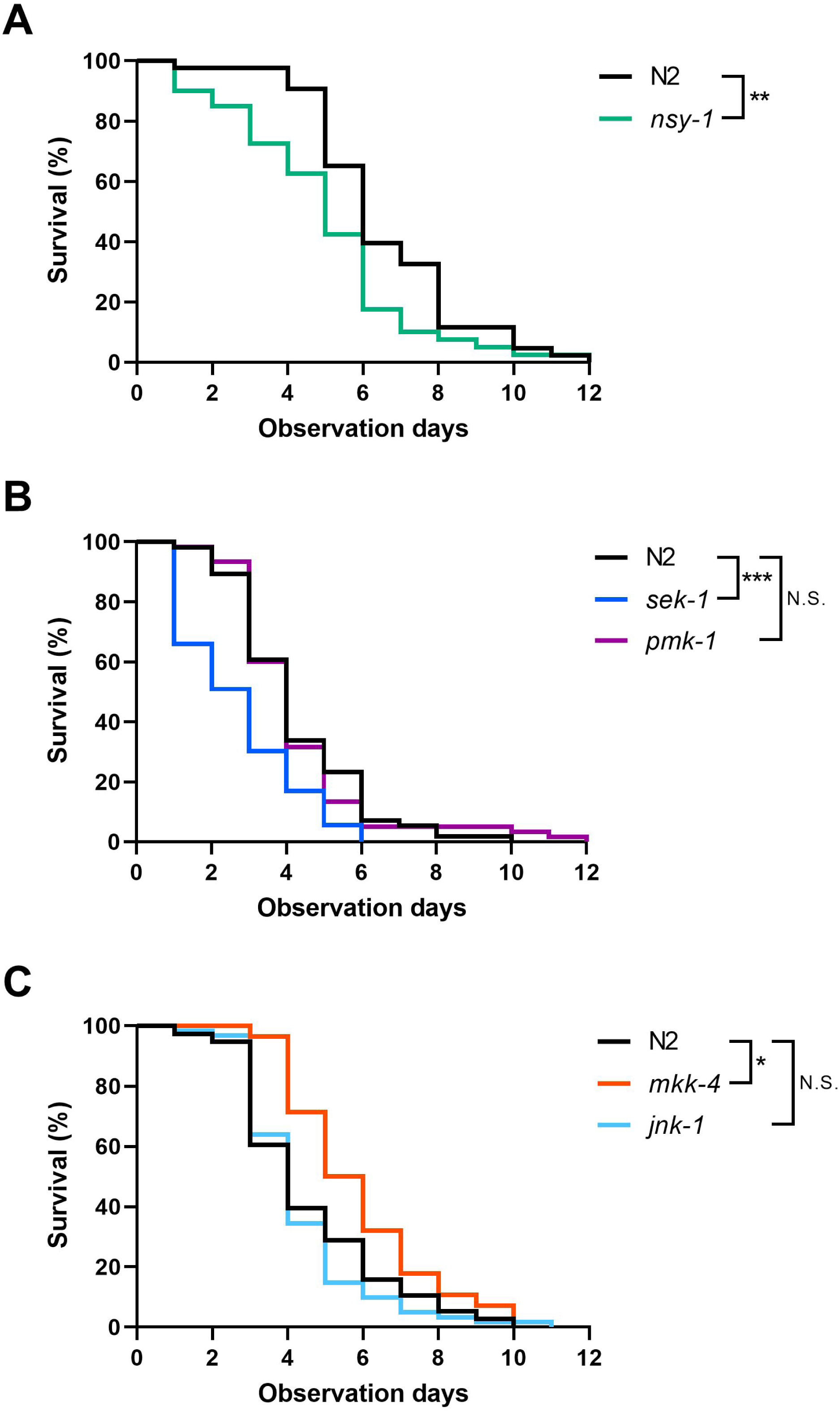
Lifespan assay of *Caenorhabditis elegans* loss-of-function mutants fed *Malassezia furfur*. (A) Survival curves of N2 (wild-type) and *nsy-1* loss-of-function mutant worms fed *M. furfur* (N2, n = 43; *nsy-1*, n = 40). (B) Survival curves of N2 and p38 pathway loss-of-function mutants (*sek-1* and *pmk-1*) worms fed *M. furfur* (N2, n = 56; *sek-1*, n = 53; *pmk-1*, n = 60). (C) Survival curves of N2 and c-Jun N-Terminal Kinase (JNK) pathway loss-of-function mutants (*mkk-4* and *jnk-1*) worms fed *M. furfur* (N2, n = 38; *mkk-4*, n = 28; *jnk-1*, n = 61). For (A) through (C), young adult worms (3-d-old) represent day 0 of observation. Survival rates were calculated using the Kaplan–Meier method and asterisks indicate a significant difference compared to N2 worms (control) using the log-rank (Mantel-Cox) test. ***, P < 0.001; **, P < 0.01; *, P < 0.05; N.S., not significant.

### Effect of *M. furfur* infection on *C. elegans* genes

RNA sequencing was performed to comprehensively investigate the genes, the expression of which was affected by *M. furfur* infection (**Table S1**). Gene Ontology (GO) enrichment terms were determined for genes upregulated or downregulated by >4-fold following *M. furfur* infection. For genes with elevated expression in the *M. furfur*-fed group, GO terms for biological processes (BP) associated with biological defense, such as “response to biotic stimulus,” were significantly enriched (**Table S2**). The downregulated genes in the *M. furfur*-fed group were similarly enriched in GO terms associated with biological defense, such as “immune system process” (**Table S3**). GO analysis was performed for genes, the expression of which increased in the *M. furfur*-fed N2 group compared to that in the OP-fed N2 group, and mRNA expression levels were quantified using real-time PCR. Of the 22 upregulated genes associated with biological defense, five were excluded because they could not be evaluated using real-time PCR. The PCR results showed significant increase in the expression of three genes encoding C-type lectin domain-containing proteins (**Fig. 3A**) and three encoding antimicrobial peptides (**Fig. 3B**) in *M. furfur*-fed N2 group. Additionally, of the seven genes categorized into other functions, six (*C50F4.9*, *cht-1*, *cyp-37B1*, *endu-2*, *fat-3*, and *math-38*) were significantly upregulated in the *M. furfur*-fed N2 group, whereas one (*sodh-1*) was not (**Fig. 3C**). Furthermore, four genes (*C14C11.4*, *C18D11.6*, *F52E1.5*, and *F53A9.8*) with unknown detailed functions were also upregulated (**Fig. 3D**).

**Fig 3.**
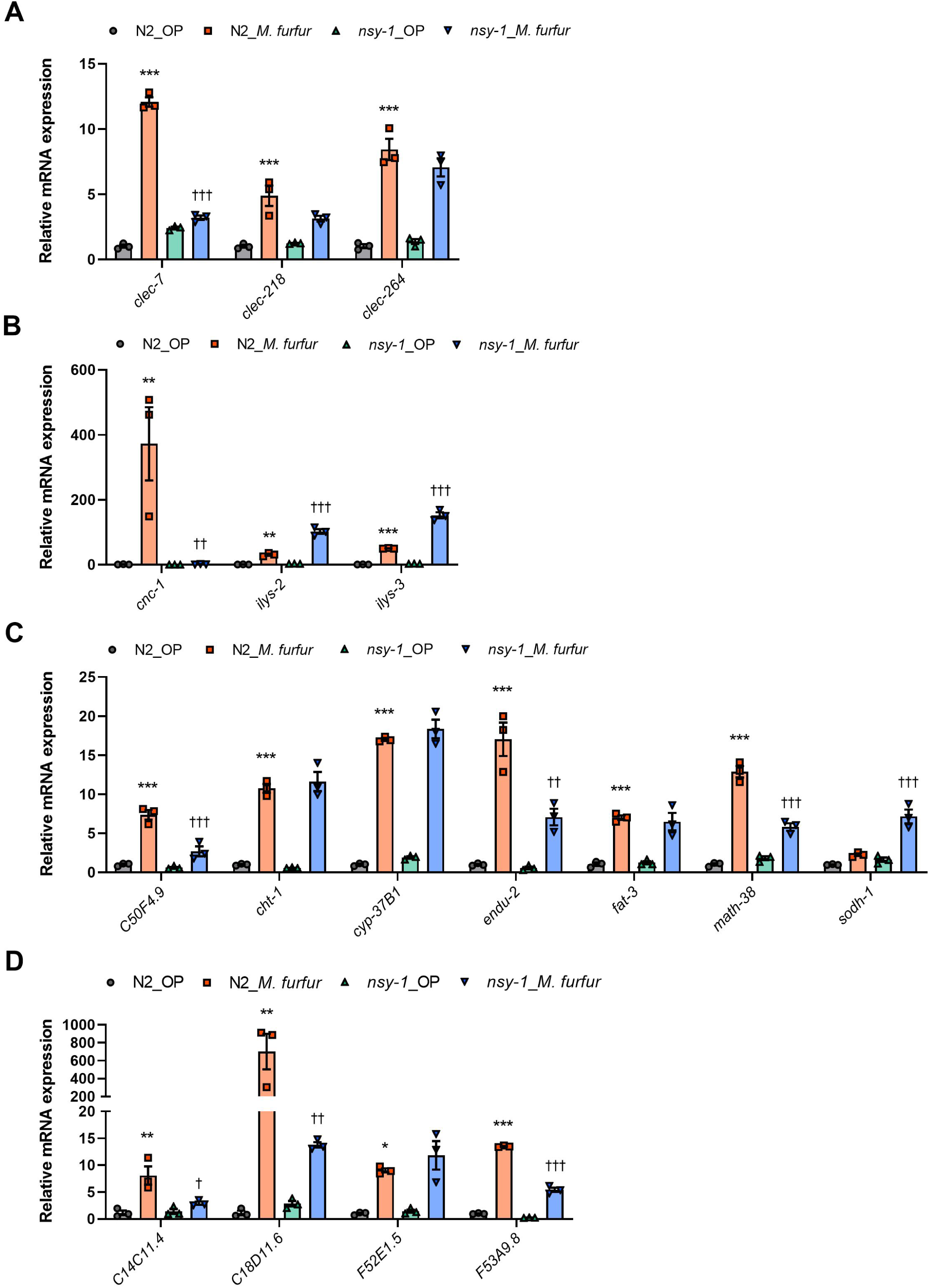
Expression of genes related to top-ranked Gene Ontology terms that were enriched in the N2 (wild-type) *Malassezia furfur*-fed *Caenorhabditis elegans* group. Genes related to C-type lectin domain-containing protein (A), antimicrobial peptide (B), other functions in response to biotic stimulus excluding those for C-type lectin or antimicrobial peptide (C), and unknown detailed function (D) are shown. For (A) through (D), real-time PCR was used to determine mRNA expression levels of genes in N2 and *nsy-1* loss-of-function mutants fed *Escherichia coli* OP50 (OP) or *M. furfur* relative to the control group (N2_OP). Data are presented as means ± standard error of mean (SEM) of three independent experiments, each normalized to three reference genes (*act-1*, *cyc-1*, or *tba-1*). Asterisks indicate statistically significant differences between N2_OP and N2_*M. furfur* groups. ***, P < 0.001; **, P < 0.01; *, P < 0.05. Daggers indicate statistically significant differences between N2_*M. furfur* and *nsy-1*_*M. furfur* groups. †††, P < 0.001; ††, P < 0.01; †, P < 0.05.

Real-time PCR was performed to determine whether specific genes in the *nsy-1* loss-of-function mutant are involved in the immune response to *M. furfur*. The results showed that the expression of eight genes (*clec-7*, *cnc-1*, *C50F4.9*, *endu-2*, *math-38*, *C14C11.4*, *C18D11.6*, and *F53A9.8*) was significantly suppressed in the *M. furfur*-fed *nsy-1* loss-of-function mutant group compared to that in the *M. furfur*-fed N2 group. In contrast, the expression of *ilys-2*, *ilys-3*, and *sodh-1* was significantly increased in the *nsy-1* loss-of-function mutant group. No significant differences were observed in the expression of six genes (*clec-218*, *clec-264*, *cht-1*, *cyp-37B1*, *fat-3*, and *F52E1.5*) between the *M. furfur*-fed N2 group and *M. furfur*-fed *nsy-1* loss-of-function mutant group (**Fig. 3A through D**).

### Effect of lactic acid bacteria and *M. furfur* mixture feeding on the lifespan of *C. elegans*

We examined whether feeding lactic acid bacteria, known for their effects on longevity and resistance to *Salmonella* infection in *C. elegans* (22), could improve the lifespan of worms shortened by *M. furfur* pathogenicity. Lifespan assays showed that N2 worms fed a 1:1 mixture of *Lacticaseibacillus rhamnosus* (LR) or *Lacticaseibacillus helveticus* (LH) and *M. furfur* had significantly longer lifespans than those fed OP + *M. furfur* (**Fig. 4A and B**). In contrast, N2 worms fed a 1:1 mixture of *Lactiplantibacillus plantarum* (LP) and *M. furfur* did not show improved survival rates (**Fig. 4B**).

**Fig 4.**
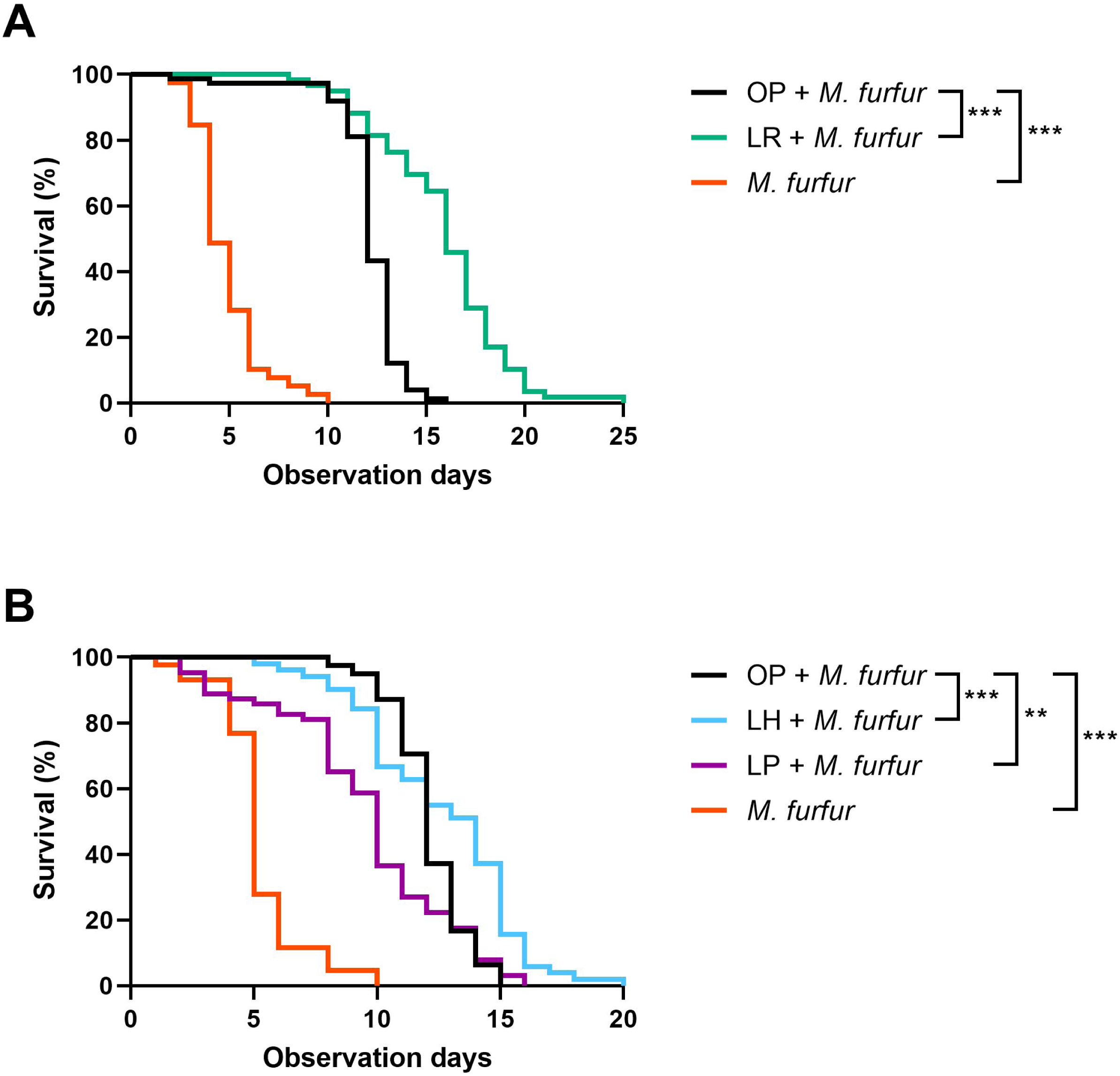
Effects of lactic acid bacteria on the lifespan of *Caenorhabditis elegans* fed *Malassezia furfur.* (A) Survival curves of N2 (wild-type) worms fed with a 1:1 mixture of *Escherichia coli* OP50 and *M. furfur* (OP + *M. furfur* (1:1), n = 74), a 1:1 mixture of *Lacticaseibacillus rhamnosus* and *M. furfur* (LR + *M. furfur* (1:1), n = 59), or only *M. furfur* (*M. furfur*, n = 39). (B) Survival curves of N2 worms fed a 1:1 mixture of OP and *M. furfur* (OP + *M. furfur* (1:1), n = 78), a 1:1 mixture of *Lactobacillus helveticus* and *M. furfur* (LH + *M. furfur* (1:1), n = 51), a 1:1 mixture of *Lactiplantibacillus plantarum* and *M. furfur* (LP + *M. furfur* (1:1), n = 63), or only *M. furfur* (*M. furfur*, n = 43). For (A) and (B), young adult worms (3-d-old) represent day 0 of observation. Survival rates were calculated using the Kaplan–Meier method and asterisks indicate a significant difference compared to OP + *M. furfur*-fed worms, using the log-rank (Mantel–Cox) test. ***, P < 0.001; **, P < 0.01.

### Effect of *L. rhamnosus* on *M. furfur* pathogenicity

Among the tested lactic acid bacteria, LR showed the greatest improvement in the survival rate of worms infected with *M. furfur*; therefore, the lifespan assay was conducted with an increased proportion of *M. furfur* (LR + *M. furfur* (1:4)). Even with a higher proportion of *M. furfur*, the survival rate of worms in the LR + *M. furfur* (1:4) group was better than that of those in the OP + *M. furfur* (1:4) group (**Fig. 5A and B**). We also examined the intestinal lumen of *C. elegans* fed a mixture of OP or LR and *M. furfur*. Worms fed a 1:4 mixture of OP or LR and *M. furfur*, as well as those fed only *M. furfur*, showed intestinal distensions near the pharynx the following day of feeding compared to worms fed only OP. In addition, *M. furfur* accumulation was observed in the intestines of worms in the OP + *M. furfur* and *M. furfur* groups. In the worms in the LR + *M. furfur* group, both LR and *M. furfur* were detected in the intestines (**Fig. S3**). To evaluate the effect of LR on *M. furfur* pathogenicity, changes in body size and intestinal barrier damage in worms fed a 1:4 mixture of LR and *M. furfur* were assessed. Projection areas were measured in worms beginning at 4 d of age (the day after grouping) to 6 d of age. The body area of worms did not differ significantly between the OP + *M. furfur* and LR + *M. furfur* groups (**Fig. 5C**). The Smurf assay was performed on worms of age 6, 10, and 13 d. At 6 d of age, nematodes in the *M. furfur* group showed more body cavity leakage than those in the other groups, while no significant differences were observed among the OP, OP + *M. furfur*, and LR + *M. furfur* groups (**Fig. 5D**). At 10 d of age, nematodes in the LR + *M. furfur* group showed significantly lower body-cavity leakage than those in the OP + *M. furfur* group. Additionally, at 13 d of age, worms in the LR + *M. furfur* group displayed fewer dye leaks into the body cavity than those in the OP and OP + *M. furfur* groups (**Fig. 5E and F**).

**Fig 5.**
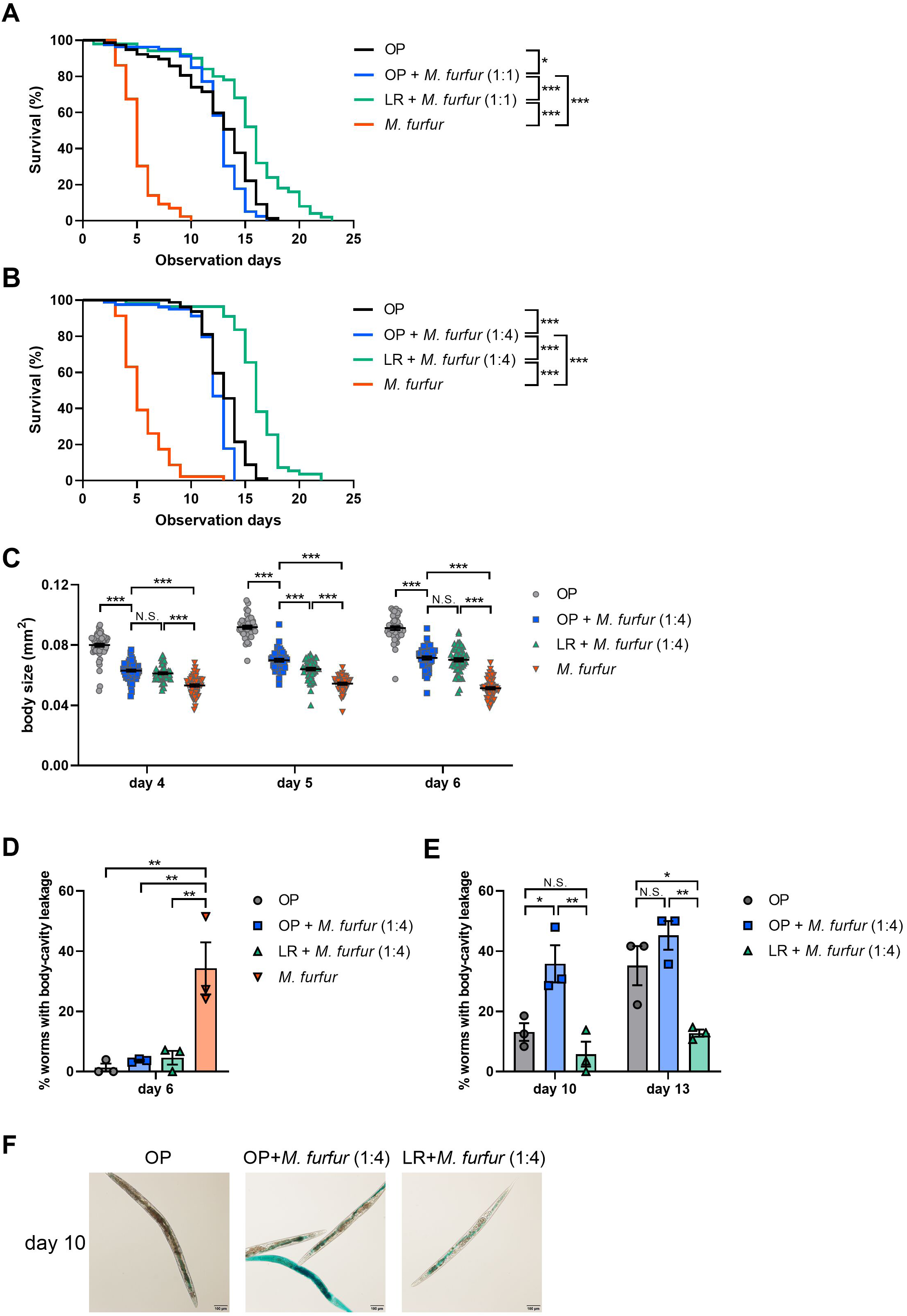
Evaluation of lactic acid bacteria protection against *Malassezia furfur* virulence in *Caenorhabditis elegans*. (A) Survival curves of N2 (wild-type) worms fed only *Escherichia coli* OP50 (OP, n = 77), a 1:1 mixture of OP and *M. furfur* (OP + *M. furfur* (1:1), n = 79), a 1:1 mixture of *Lacticaseibacillus rhamnosus* and *M. furfur* (LR + *M. furfur* (1:1), n = 50), or only *M. furfur* (*M. furfur*, n = 43). (B) Survival curves of N2 worms fed a mixture with a high proportion of *M. furfur* (OP, n = 79; OP + M. furfur (1:4), n = 79; LR + *M. furfur* (1:4), n = 55; *M. furfur*, n = 46). For (A) and (B), young adult worms (3-d-old) represent day 0 of observation. Survival rates were calculated using the Kaplan–Meier method and asterisks indicate a significant difference compared to OP + *M. furfur*-fed worms using the log-rank (Mantel–Cox) test. ***, P < 0.001; *, P < 0.05. (C) Body size of worms fed OP, OP + *M. furfur* (1:4), LR + *M. furfur* (1:4), or only *M. furfur* from 4 to 6 d of age. Asterisks indicate significant differences compared to OP + *M. furfur* (1:4)-fed worms. (D) The percentage of body-cavity leakage worms fed OP, OP + *M. furfur* (1:4), or LR + *M. furfur* (1:4) was compared to those fed *M. furfur* at 6 d of age. (E) The percentage of worms with body-cavity leakage fed OP and LR + *M. furfur* (1:4) was compared to those fed OP + *M. furfur* (1:4) at 10 and 13 d of age. For (C) through (E), results are shown as individual plots and means ± standard error of mean (SEM). Statistical analysis was performed using one-way ANOVA and Tukey’s multiple comparison tests. ***, P < 0.001; **, P < 0.01; *, P < 0.05; N.S., not significant. (F) Representative images of 10-d-old worms fed OP, OP + *M. furfur* (1:4), or LR + *M. furfur* (1:4) and stained with blue dye. Scale bar, 100 µm.

## Discussion

This study was conducted to characterize *M. furfur* pathogenicity and evaluate the protective effects of probiotics, specifically lactic acid bacteria, against it, using *C. elegans* as a model host. We first assessed the impact of *M. furfur* infection on nematodes to evaluate its virulence. *M. furfur*-fed *C. elegans* exhibited significantly shorter lifespans than the controls, indicating that *M. furfur* is pathogenic to nematodes. Additionally, HK-*M. furfur* also shortened the lifespan of nematodes, although to a lesser extent than live *M. furfur* did. Heat-killed *Cryptococcus neoformans*, the yeast that causes cryptococcosis, is pathogenic to nematodes, suggesting that part of the pathogenicity of this yeast may be due to cell wall fragments or toxins (26). In the present study, compared to OP-feeding, both *M. furfur* and HK-*M. furfur* infection significantly reduced the body surface area of *C. elegans*. *Malassezia* contains pathogenic factors, such as galactomannans in the cell wall, and various indole compounds, including malassezin (a tryptophan metabolite) (9, 27, 28). Aryl hydrocarbons are heat-stable; therefore, part of *M. furfur*’s pathogenicity could be attributed to cellular components (CC), such as metabolites and cell wall of the fungus, which are less affected by heating. Furthermore, *C. albicans* mutants that have a reduced ability to form mycelia are significantly less toxic to nematodes(29). Similar to *C. albicans*, *M. furfur* can exist in both yeast and hyphal forms(30, 31); therefore, mycelial formation may also play a significant role in the pathogenicity of *M. furfur*. In the present study, several nematodes fed live *M. furfur* exhibited dye leakage, suggesting that *M. furfur* damages the intestinal barrier. *C. elegans* lacks acquired immunity; therefore, the intestinal barrier is an important defense mechanism against pathogenic microorganisms. A similar disruption is caused by enteropathogenic *E. coli* in *C. elegans* (32). In contrast, HK-*M. furfur* did not cause significant intestinal damage, suggesting that virulence factors of dead *M. furfur* do not include components responsible for intestinal destruction.

*M. furfur-*fed *nsy-1* and *sek-1* loss-of-function mutant *C. elegans* showed a decrease in lifespan, while the *pmk-1* and *jnk-1* loss-of-function mutants showed no change in lifespan. Notably, as compared to that of N2, the lifespan of the *mkk-4* loss-of-function mutant was extended after *M. furfur* infection, suggesting that this mutant may be detrimental to *M. furfur*. NSY-1 is a homolog of the mammalian MAPKKK ASK1 that activates JNK and p38 MAPKs(33), and the p38 MAP kinase pathway consists of NSY-1-SEK-1-PMK-1 (33). Mammalian MKK4 MAPKK activates both p38 and JNK MAPKs,(34) and MKK-4 and JNK-1 in *C. elegans* encode homologs of mammalian MKK4 and JNK, respectively (34). These results suggest that *nsy-1* and *sek-1* play important roles in the p38 MAPK-related response to *M. furfur* pathogenicity. Insulin-like signaling pathways, transforming growth factor-β (TGF-β)-like pathways, and programmed cell death pathways are involved in the response to various pathogen infections in *C. elegans*; these pathways may interact with the MAPK pathway (35–40).

GO analysis revealed that among the genes, the expression of which increased or decreased more than 4-fold following *M. furfur* infection, a substantial number were involved in defense responses, suggesting that the expression of genes involved in defense responses is regulated by *M. furfur* infection. Further GO analysis revealed that the genes, the expression of which were upregulated by *M. furfur* infection, were involved in the defense response to gram-positive bacteria. This finding suggests that defense responses mechanisms against gram-positive bacteria and fungi may be partially similar in *C. elegans*. In contrast, *M. furfur* infection downregulated the expression of genes involved in the immune system; however, we only focused on genes upregulated by *M. furfur* infection. Real-time PCR of N2 nematodes demonstrated that the expression of *clec-7*, *clec-218*, and *clec-264,* which encode C-type lectin domain-containing proteins involved in pathogen recognition, along with *cnc-1*, *ilys-2*, and *ilys-3*, which encode antimicrobial peptides, increased with *M. furfur* infection. C-type lectin-like proteins are involved not only in recognition but also in pathogen elimination as soluble secretory proteins (41). Some caenacin genes, including *cnc-1* of *C. elegans*, are induced by infection with the fungus *Drechmeria coniospora* (39); furthermore, the invertebrate lysozyme *ilys-3* is upregulated upon infection with *C. albicans* (42). These results suggest that the expression of multiple genes associated with biological defense is activated in *M. furfur* infection. Additionally, *cht-1*, one of the genes, the expression of which was upregulated in *M. furfur* infection, encodes an endochitinase and is involved in *C. albicans* infection (21). In contrast, *sodh-1*, which encodes an alcohol dehydrogenase and has been implicated in the fungus *Purpureocillium lavendulum* infection (43), did not show increased expression upon *M. furfur* infection. To further investigate the mechanism of the immune response, we compared the gene expression levels in the *nsy-1* loss-of-function mutant *C. elegans* with those in N2 nematodes. Of the 16 genes that showed significant upregulated expression upon *M. furfur* infection in N2, the expression of eight was significantly suppressed in the *nsy-1* loss-of-function mutant nematodes exposed to *M. furfur*, suggesting the dependence of these genes on NSY-1 for their activation in response to *M*. *furfur*. In contrast, *ilys-2*, *ilys-3*, and *sodh-1* were upregulated in the *nsy-1* loss-of-function mutant. The expression of *ilys-3* is upregulated in an extracellular signal-regulated MAP kinase (ERK)-MAPK pathway-dependent manner (42). The loss of *nsy-1* may have triggered the activation of other immune response pathways to compensate, leading to increased expression of *ilys -2*, *ilys-3*, and *sodh-1* downstream of these pathways.

*Malassezia* is associated with inflammatory bowel disease (10). In the present study, *M. furfur* caused intestinal barrier disruption in *C. elegans*; therefore, we investigated the protective effect of probiotic bacteria in *M. furfur* infection. The results of mixed feeding with each of the three lactic acid bacterial species (LR, LH, or LP) and *M. furfur* showed that the survival rates of the nematodes were higher in the LR or LH and *M. furfur* mixed feeding group than the OP + *M. furfur* group, suggesting that LR and LH protect nematodes from the pathogenic effects of *M. furfur*. The OP + *M. furfur* mixed feeding group also showed an increased survival rate compared to the *M. furfur* infection group, suggesting that OP may also mitigate *M. furfur* virulence, though to a lesser extent than LR. In contrast, the LP + *M. furfur* group did not extend the survival rate of the OP + *M. furfur* group, indicating that different types of lactic acid bacteria offer varying degrees of protection against *M. furfur* pathogenicity. Feeding with LR, LH, or LP extends the lifespan of nematodes compared to feeding with OP (22), suggesting that lifespan extension and protection against pathogens may be mediated through different mechanisms. For instance, wild-type nematodes pretreated with the isolate *Lacticaseibacillus zeae* before infection with enterotoxigenic *Escherichia coli* (ETEC) showed a significantly extended lifespan compared to those pretreated with OP before infection (44).

Among the three species of lactic acid bacteria, LR provided the most significant improvement in *C. elegans* survival against *M. furfur*; therefore, we evaluated whether feeding LR could alleviate the reduction in nematode body area and intestinal barrier damage caused by *M. furfur*. No significant differences in body area were observed between the OP + *M. furfur* and LR + *M. furfur* groups on the last day of body area measurements, suggesting that although mixed LR feeding did slightly slow the growth rate of nematodes compared to mixed OP feeding, it did not significantly alter the final body size. *Lactobacillus* retention and intestinal distention were also observed in *C. elegans* fed LR + *M. furfur*, indicating that LR does not eliminate the intestinal distention caused by *M. furfur*. *Enterococcus faecalis* causes intestinal distention and shortens survival, while *E. faecium* causes intestinal distention without affecting the lifespan of *C. elegans* (45), suggesting that intestinal distention and the accumulation of bacteria do not necessarily affect the lifespan of *C. elegans*. It is possible that LR prevented *M. furfur* from contacting the host intestinal tract by colonizing the intestinal tract, thereby preventing the induction of an excessive immune response. Intestinal staining results showed that significantly more 6-d-old nematodes in the *M. furfur* group exhibited dye leakage than those in the OP, OP + *M. furfur*, and LR + *M. furfur* groups, suggesting that mixed feeding with OP or LR could prevent damage to the nematode intestinal barrier caused by *M. furfur*. Moreover, significantly fewer 10- and 13-d-old nematodes in the LR + *M. furfur* group showed dye leakage than those in the OP + *M. furfur* group, suggesting that LR protects the intestinal barrier more effectively than OP. *Lacticaseibacillus zeae* regulates nematode signaling and the production of antimicrobial peptides and other defense effectors during ETEC infection (44). The LR strain Lcr35 significantly inhibited *C. albicans* growth and adhesion to Caco-2 cells in vitro (46). The protective mechanism of LR in *M. furfur* infection in nematodes may include direct antimicrobial effects against *M. furfur* via antimicrobial peptides, and/or indirect effects via activation of host immunity, such as strengthening the intestinal barrier and enhancing intestinal repair.

In conclusion, this study characterized the virulence of *M. furfur* and identified the protective effect of LR against *M. furfur* infection in *C. elegans* (**Fig. 6A**). The shortened lifespan of *C. elegans* infected with *M. furfur* was linked to the p38 signaling pathways specifically involving *nsy-1* and *sek-1* (**Fig. 6B**). Although *pmk-1* has been reported to be a factor that acts downstream of *sek-1* (33), *pmk-1* was not involved in *M. furfur* infection, suggesting that there may be another unknown pathway. Gene expression analyses in *C. elegans* exposed to *M. furfur* showed elevated expression levels for genes involved in host defense, C-type lectin domain-containing proteins, and antimicrobial peptides. We believe that our findings will lead to a better understanding of the diseases associated with *M. furfur* and provide insights into developing novel preventive and therapeutic approaches using probiotics.

**Fig 6.**
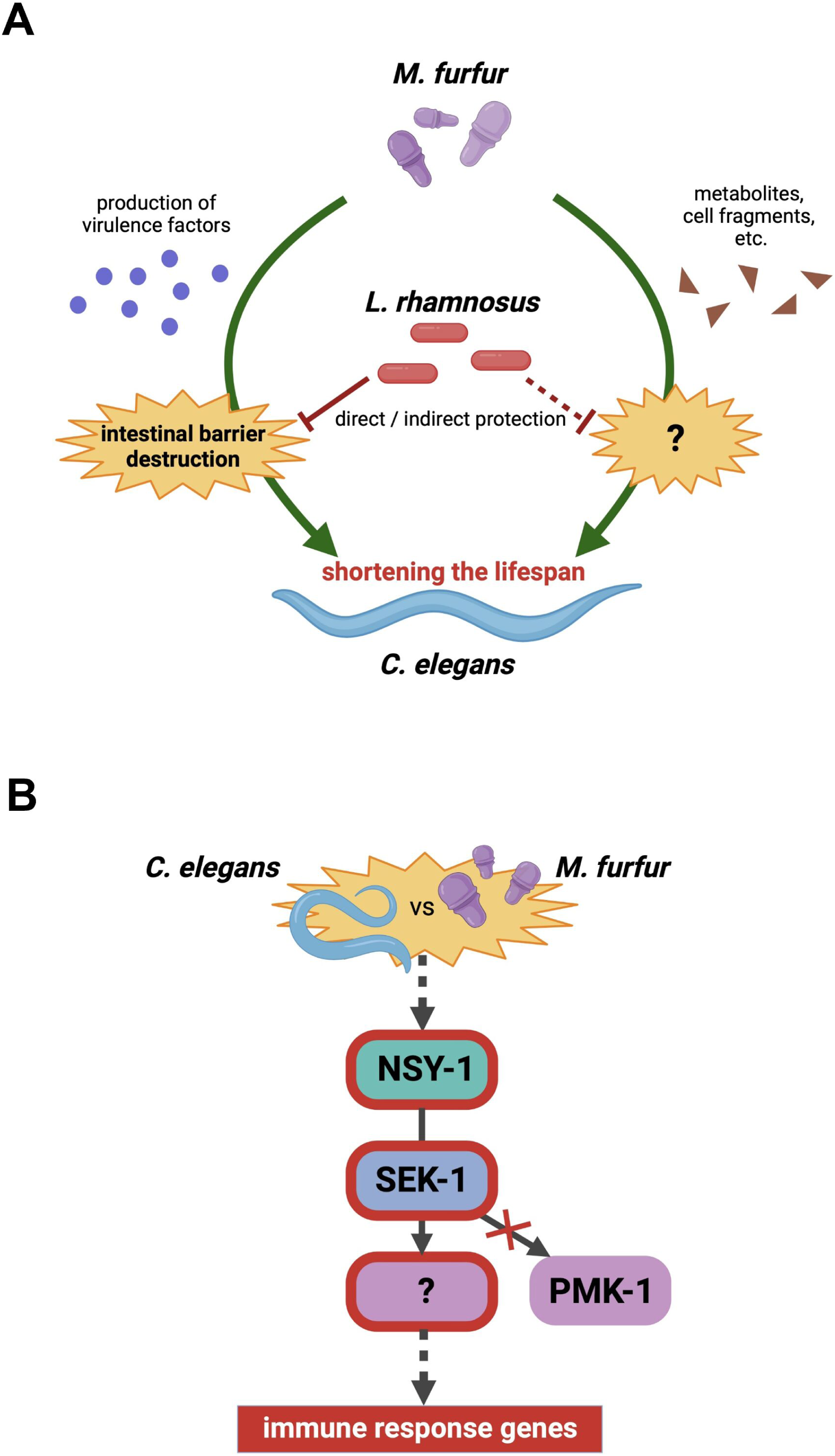
Schematic representation of the pathogenicity of *Malassezia furfur* in *Caenorhabditis elegans*. (A) The protective effect of *Lacticaseibacillus rhamnosus*. (B) The signaling pathway.

## Materials and Methods

### Bacterial strains and culture conditions

*Malassezia furfur* NBRC0656, *Lacticaseibacillus rhamnosus* NBRC14710, *Lactobacillus helveticus* NBRC15019, and *Lactiplantibacillus plantarum* NBRC15891 were sourced from the Biological Resource Center of the National Institute of Technology and Evaluation (NBRC, Tokyo, Japan). *Malassezia furfur* was cultured in CHROMagar Malassezia/Candida media (Kanto Chemical, Tokyo, Japan) at 30 °C for 72 h. Lactic acid bacteria were cultivated anaerobically in No. 804 medium (LR and LH), and in MRS medium (*L. plantarum;* Kanto Chemical). No. 804 medium was prepared by mixing 5.0 g/L Bacto peptone (Thermo Fisher Scientific, Waltham, MA, USA), 5.0 g/L yeast extract (Thermo Fisher Scientific), 5.0 g/L glucose (Fujifilm Wako Pure Chemical, Osaka, Japan), and 1.0 g/L MgSO_4_·7H_2_O (Nacalai tesque, Kyoto, Japan), and 20 g/L agar (Fujifilm Wako Pure Chemical) was added as needed. *Escherichia coli* OP50 was grown on tryptone soya agar (Shimadzu Diagnostics, Tokyo, Japan) at 37 °C for 24 h and used as *C. elegans* standard feed. The bacterial/fungal lawns were suspended in M9 buffer (5 mM potassium phosphate, 1 mM CaCl_2_, and 1 mM MgSO_4_) at 10 mg/50 µL to feed *C. elegans*. Heat-killed organisms were prepared by incubation at 100 °C for 15 min. The bacterial/fungal suspension (50 μL) was spread on 5-cm peptone-free-modified nematode growth media (mNGM; 1.7% w/v agar, 50 mM NaCl, 1 mM CaCl_2_, 5 μg/mL cholesterol, 25 mM KH_2_PO_4_, and 1 mM MgSO_4_) plates.

### Nematodes and growth conditions

*C. elegans* Bristol strain N2 (wild-type) and its loss-of-function mutant strains AU3 *nsy-1* (*ag3*), KU4 *sek-1* (*km4*), KU25 *pmk-1* (*km25*), CZ4213 *mkk-4* (*ju91*), and VC8 *jnk-1* (*gk7*) were provided by the Caenorhabditis Genetics Center, University of Minnesota (Minneapolis, MN, USA). The nematodes were maintained and propagated on nematode growth media following an established method (47). Eggs were obtained by exposing mature *C. elegans* to a solution containing sodium hypochlorite and sodium hydroxide. The eggs were cultivated in M9 buffer at 25 °C for 1 d to promote hatching and synchronization. Synchronized larval stage 1 worms were collected via centrifugation, transferred onto fresh mNGM plates spread with OP, and incubated at 25 °C for 2 d until they reached the young adult (3-d-old) stage. To prevent contamination by progeny and dead worms due to internal hatching, 3-d-old *C. elegans* were treated with 2’-deoxy-5-fluorouridine (Tokyo Chemical Industry, Tokyo, Japan) for all experiments (48).

### Lifespan assay

Synchronized 3-d-old N2 or loss-of-function mutant worms were placed on 5-cm mNGM plates (40 animals per plate) already spread with bacteria and/or fungi and incubated at 25 °C. Worms were transferred daily to fresh plates for the first 4 d and every other day thereafter. The number of live and dead worms was recorded daily. A worm was considered dead if it failed to respond to gentle touch with a worm picker. Worms that crawled into the agar or died by adhering to the plate wall were excluded from the analyses. Each assay was performed in duplicate. Worm survival curves were calculated using the Kaplan–Meier method. Mean life span (MLS) was estimated using the following equation (49):

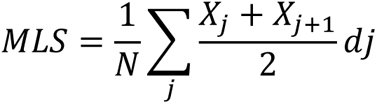

where *N* is the total number of worms and *d_j_* is the number of worms that died in the age interval (*X_j_* to *X_j_*_+1_). The standard error (SE) of the estimated mean lifespan was calculated using the following equation:

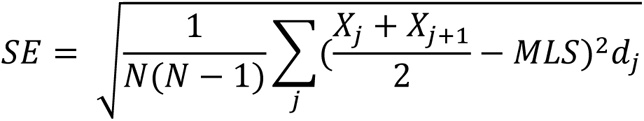

Data of MLS ± SE for all experiments in this study are provided in **Table S4**.

### Measurement of body size

Three-day-old adult worms were placed on mNGM plates with bacteria and/or fungi and incubated at 25 °C. The size of live worms was measured every 24 h until they reached 6 d of age. Images of mature nematodes were captured and quantified using an all-in-one fluorescence microscope (BZ-X800, Keyence, Osaka, Japan) and a hybrid cell count application (BZ-H3C, Keyence) in the BZ-X Analyzer software (BZ-H3A, Keyence). The worm projection area was used as the body size index. The assay was performed with >40 worms per group for each time point.

### Intestinal barrier function assay (Smurf assay)

The integrity of the intestinal barrier function was examined according to a previously described method (32), with modifications. Three-day-old adult worms were placed on mNGM plates containing the feed described above and incubated at 25 °C. The worms were removed from the mNGM plates at 4–6 d of age or at 6, 10, and 13 d of age and further cultured in liquid media containing HK-OP mixed with blue food dye (Acid Blue 9; Tokyo Chemical Industry, 5% [wt/vol] in NGM liquid solution) at 25 °C for 3 h. The worms were washed in M9 buffer until the blue color of the dye was not visible. Then, the worms were observed on a slide with 50 mM sodium azide for the presence or absence of blue food dye in the body cavity using an upright microscope (BX53, Olympus, Tokyo, Japan) and a digital camera (DP73, Olympus) at x100 magnification. The stained worms were counted to determine body cavity leakage. Three independent experiments were performed with >15 worms per group for each time point.

### RNA extraction and sequencing

Three-day-old worms were placed on mNGM plates coated with OP or *M. furfur* and incubated at 25 °C for 6 h. Approximately 200 worms per group were collected, washed at least three times with M9 buffer containing 0.2% gelatin, and stored in RNAprotect Tissue Reagent (Qiagen, Hilden, Germany) at −80 °C until RNA extraction. Then, the thawed nematode suspensions were ground using a microtube pestle (Scientific Specialties, Lodi, CA, USA) and total RNA was extracted using TRIzol reagent (Thermo Fisher Scientific) and the RNeasy Mini Kit (Qiagen). RNA sequencing was performed by DNAFORM (Yokohama, Japan) as previously described (50). The read count of gene features was performed using the feature Counts tool (version 1.6.1) (51), following which quantitative differential expression analysis was conducted between the control (N2_OP) and test-fed groups (N2_*M. furfur*) using DESeq2 (version 1.30.0) (52). Enrichment analysis based on the GO categories for BP, molecular functions, and CC was performed using clusterProfiler (version 3.6.0) (53) on genes that exhibited a |log_2_FC| of ≥2 and a base mean of ≥5 (**Tables S2 and S3**).

### Reverse transcription and real-time PCR

cDNA was generated using the QuantiTect Reverse Transcription Kit (Qiagen). Real-time quantitative PCR was conducted using the StepOnePlus Real-Time PCR Systems (Thermo Fisher Scientific) with the Power SYBR Green PCR Master Mix (Thermo Fisher Scientific) under the following thermal cycling conditions: initial denaturation at 95 °C for 10 min, followed by 40 cycles of denaturation at 95 °C for 15 s and annealing/extension at 60 °C for 1 min. Samples from three biological replicates were analyzed. Relative mRNA expression was calculated using the ΔΔCt method (54) and normalized to the expression of three reference genes: *act-1*, *tba-1*, and *cyc-1*. The primers used for real-time PCR are listed in **Table S5**.

### Statistical analyses

Statistical analyses were performed using the GraphPad Prism 8 software (GraphPad Prism Software, San Diego, CA, USA). The survival of *C. elegans* was calculated using the Kaplan–Meier method, and statistical significance was assessed using the log-rank (Mantel-Cox) test. Body size, intestinal barrier function, and relative mRNA expression were compared using one-way ANOVA and Tukey’s multiple comparison test.

## Data availability

Data supporting the findings of this study are available from the Gene Expression Omnibus (GEO) (GSE260976).

## Acknowledgments

This study was supported by JSPS KAKENHI grant number 22K1975. We would like to acknowledge the *Caenorhabditis* Genetics Center (University of Minnesota, Minneapolis, MN, supported by the National Institutes of Health National Center for Research Resources) and the National Institute of Genetics through the National BioResource Project of MEXT, Japan, for providing the *C. elegans* strains, and the National Institute of Technology and Evaluation (NITE) Biological Resource Center for providing the *Malassezia furfur*, *Lacticaseibacillus rhamnosus*, *Lactobacillus helveticus*, and *Lactiplantibacillus plantarum* strains.

## Author contribution

C. K. and E. K. N. designed the study. C.K., A.T., X.W., and Y.T. conducted molecular and genetic experiments, survival assays, and data analyses. C.K., Y.T., and E.K.N. drafted the manuscript. All the authors have read and approved the manuscript.

## References

1. Gao Z, Perez-Perez GI, Chen Y, Blaser MJ. 2010. Quantitation of major human cutaneous bacterial and fungal populations. J Clin Microbiol 48:3575–3581.

2. Prohic A, Jovovic Sadikovic T, Krupalija-Fazlic M, Kuskunovic-Vlahovljak S. 2016. *Malassezia* species in healthy skin and in dermatological conditions. Int J Dermatol 55:494–504.

3. Borgers M, Cauwenbergh G, Van De Ven M -A, Hernanz AP, Degreef H. 1987. Pityriasis versicolor and Pityrosporum ovale. Morphogenetic and ultrastructural considerations. Int J Dermatol 26:586–589.

4. Elewski B. 1990. Does Pityrosporum ovale have a role in psoriasis? Arch Dermatol 126:1111– 1112.

5. Piérard-Franchimont C, Arrese JE, Piérard GE. 1995. Immunohistochemical aspects of the link between *Malassezia ovalis* and seborrheic dermatitis. Journal of the European Academy of Dermatology and Venereology 4:14–19.

6. Ashbee HR, Leck AK, Puntis JWL, Parsons WJ, Evans EG V. 2002. Skin colonization by *Malassezia* in neonates and infants. Infect Control Hosp Epidemiol 23:212–216.

7. Prohic A, Ozegovic L. 2007. *Malassezia* species isolated from lesional and non-lesional skin in patients with pityriasis versicolor. Mycoses 50:58–63.

8. Patĩo-Uzcátegui A, Amado Y, Cepero De García M, Chaves D, Tabima J, Motta A, Cárdenas M, Bernal A, Restrepo S, Celis A. 2011. Virulence gene expression in *Malassezia* spp from individuals with seborrheic dermatitis. J Invest Dermatol 131:2134–2136.

9. Gaitanis G, Magiatis P, Stathopoulou K, Bassukas ID, Alexopoulos EC, Velegraki A, Skaltsounis AL. 2008. AhR ligands, malassezin, and indolo[3,2-b]carbazole are selectively produced by *Malassezia furfur* strains isolated from seborrheic dermatitis. J Invest Dermatol 128:1620–1625.

10. Limon JJ, Tang J, Li D, Wolf AJ, Michelsen KS, Funari V, Gargus M, Nguyen C, Sharma P, Maymi VI, Iliev ID, Skalski JH, Brown J, Landers C, Borneman J, Braun J, Targan SR, McGovern DPB, Underhill DM. 2019. *Malassezia* is associated with Crohn’s disease and exacerbates colitis in mouse models. Cell Host Microbe 25:377–388.e6.

11. Abdillah A, Ranque S. 2021. Chronic diseases associated with *Malassezia* yeast. J Fungi (Basel) 7.

12. Lorch JM, Palmer JM, Vanderwolf KJ, Schmidt KZ, Verant ML, Weller TJ, Blehert DS. 2018. *Malassezia vespertilionis* sp. nov.: a new cold-tolerant species of yeast isolated from bats. Persoonia 41:56–70.

13. Gaitanis G, Magiatis P, Hantschke M, Bassukas ID, Velegraki A. 2012. The *Malassezia* genus in skin and systemic diseases. Clin Microbiol Rev 25:106–141.

14. Rhimi W, Theelen B, Boekhout T, Otranto D, Cafarchia C. 2020. *Malassezia* spp. yeasts of emerging concern in fungemia. Front Cell Infect Microbiol 10.

15. Lamas B, Natividad JM, Sokol H. 2018. Aryl hydrocarbon receptor and intestinal immunity. Mucosal Immunol 11:1024–1038.

16. Tajima M, Sugita T, Nishikawa A, Tsuboi R. 2008. Molecular analysis of *Malassezia* microflora in seborrheic dermatitis patients: comparison with other diseases and healthy subjects. J Invest Dermatol 128:345–351.

17. Prohic A, Simic D, Sadikovic TJ, Krupalija-Fazlic M. 2014. Distribution of *Malassezia* species on healthy human skin in Bosnia and Herzegovina: correlation with body part, age and gender. Iran J Microbiol 6:253.

18. Finch CE, Ruvkun G. 2001. The genetics of aging. Annu Rev Genomics Hum Genet 2:435–462.

19. Nicholas HR, Hodgkin J. 2004. Responses to infection and possible recognition strategies in the innate immune system of *Caenorhabditis elegans*. Mol Immunol 41:479–493.

20. Labrousse A, Chauvet S, Couillault C, Léopold Kurz C, Ewbank JJ. 2000. *Caenorhabditis elegans* is a model host for *Salmonella typhimurium*. Curr Biol 10:1543–1545.

21. Pukkila-Worley R, Ausubel FM, Mylonakis E. 2011. *Candida albicans* infection of *Caenorhabditis elegans* induces antifungal immune defenses. PLoS Pathog 7.

22. Ikeda T, Yasui C, Hoshino K, Arikawa K, Nishikawa Y. 2007. Influence of lactic acid bacteria on longevity of *Caenorhabditis elegans* and host defense against *Salmonella enterica* serovar enteritidis. Appl Environ Microbiol 73:6404–6409.

23. Laranjeiro R, Harinath G, Hewitt JE, Hartman JH, Royal MA, Meyer JN, Vanapalli SA, Driscoll M. 2019. Swim exercise in *Caenorhabditis elegans* extends neuromuscular and gut healthspan, enhances learning ability, and protects against neurodegeneration. Proc Natl Acad Sci U S A 116:23829–23839.

24. Sakaguchi A, Matsumoto K, Hisamoto N. 2004. Roles of MAP kinase cascades in *Caenorhabditis elegans*. J Biochem 136:7–11.

25. Wang Y, Zhang L, Luo X, Wang S, Wang Y. 2017. Bisphenol A exposure triggers apoptosis via three signaling pathways in *Caenorhabditis elegans*. RSC Adv 7:32624–32631.

26. Mylonakis E, Ausubel FM, Perfect JR, Heitman J, Calderwood SB. 2002. Killing of *Caenorhabditis elegans* by *Cryptococcus neoformans* as a model of yeast pathogenesis. Proc Natl Acad Sci U S A 99:15675–15680.

27. Shibata N, Saitoh T, Tadokoro Y, Okawa Y. 2009. The cell wall galactomannan antigen from *Malassezia furfur* and *Malassezia pachydermatis* contains beta-1,6-linked linear galactofuranosyl residues and its detection has diagnostic potential. Microbiology (Reading) 155:3420–3429.

28. Krämer HJ, Podobinska M, Bartsch A, Battmann A, Thoma W, Bernd A, Kummer W, Irlinger B, Steglich W, Mayser P. 2005. Malassezin, a novel agonist of the aryl hydrocarbon receptor from the yeast *Malassezia furfur*, induces apoptosis in primary human melanocytes. Chembiochem 6:860–865.

29. Pukkila-Worley R, Peleg AY, Tampakakis E, Mylonakis E. 2009. *Candida albicans* hyphal formation and virulence assessed using a *Caenorhabditis elegans* infection model. Eukaryot Cell 8:1750–1758.

30. Gaitanis G, Velegraki A, Mayser P, Bassukas ID. 2013. Skin diseases associated with *Malassezia* yeasts: facts and controversies. Clin Dermatol 31:455–463.

31. Sudbery P, Gow N, Berman J. 2004. The distinct morphogenic states of *Candida albicans*. Trends Microbiol 12:317–324.

32. Kim J, Moon Y. 2019. Worm-based alternate assessment of probiotic intervention against gut barrier infection. Nutrients 11.

33. Sagasti A, Hisamoto N, Hyodo J, Tanaka-Hino M, Matsumoto K, Bargmann CI. 2001. The CaMKII UNC-43 activates the MAPKKK NSY-1 to execute a lateral signaling decision required for asymmetric olfactory neuron fates. Cell 105:221–232.

34. Kim DH, Feinbaum R, Alloing G, Emerson FE, Garsin DA, Inoue H, Tanaka-Hino M, Hisamoto N, Hatsumoto K, Tan MW, Ausubel FH. 2002. A conserved p38 MAP kinase pathway in *Caenorhabditis elegans* innate immunity. Science 297:623–626.

35. Garsin DA, Villanueva JM, Begun J, Kim DH, Sifri CD, Calderwood SB, Ruvkun G, Ausubel FM. 2003. Long-lived *C. elegans* daf-2 mutants are resistant to bacterial pathogens. Science 300:1921.

36. Mallo G V., Kurz CL, Couillault C, Pujol N, Granjeaud S, Kohara Y, Ewbank JJ. 2002. Inducible antibacterial defense system in *C. elegans*. Curr Biol 12:1209–1214.

37. Aballay A, Ausubel FM. 2001. Programmed cell death mediated by ced-3 and ced-4 protects *Caenorhabditis elegans* from *Salmonella typhimurium*-mediated killing. Proc Natl Acad Sci U S A 98:2735–2739.

38. Kondo M, Yanase S, Ishii T, Hartman PS, Matsumoto K, Ishii N. 2005. The p38 signal transduction pathway participates in the oxidative stress-mediated translocation of DAF-16 to *Caenorhabditis elegans* nuclei. Mech Ageing Dev 126:642–647.

39. Zugasti O, Ewbank JJ. 2009. Neuroimmune regulation of antimicrobial peptide expression by a noncanonical TGF-beta signaling pathway in *Caenorhabditis elegans* epidermis. Nat Immunol 10:249–256.

40. Aballay A, Drenkard E, Hilbun LR, Ausubel FM. 2003. *Caenorhabditis elegans* innate immune response triggered by *Salmonella enterica* requires intact LPS and is mediated by a MAPK signaling pathway. Curr Biol 13:47–52.

41. Schulenburg H, Kurz CL, Ewbank JJ. 2004. Evolution of the innate immune system: the worm perspective. Immunol Rev 198:36–58.

42. Gravato-Nobre MJ, Vaz F, Filipe S, Chalmers R, Hodgkin J. 2016. The invertebrate lysozyme effector ILYS-3 is systemically activated in response to danger signals and confers antimicrobial protection in *C. elegans*. PLoS Pathog 12.

43. Zhuang X-M, Guo Z-Y, Zhang M, Chen Y-H, Qi F-N, Wang R-Q, Zhang L, Zhao P-J, Lu C-J, Zou C-G, Ma Y-C, Xu J, Zhang K-Q, Cao Y-R, Liang L-M. 2023. Ethanol mediates the interaction between *Caenorhabditis elegans* and the nematophagous fungus *Purpureocillium lavendulum*. Microbiol Spectr 11.

44. Zhou M, Liu X, Yu H, Yin X, Nie SP, Xie MY, Chen W, Gong J. 2018. Cell signaling of *Caenorhabditis elegans* in response to enterotoxigenic *Escherichia coli* infection and *Lactobacillus zeae* protection. Front Immunol 9.

45. Yuen GJ, Ausubel FM. 2018. Both live and dead *Enterococci* activate *Caenorhabditis elegans* host defense via immune and stress pathways. Virulence 9:683–699.

46. Poupet C, Saraoui T, Veisseire P, Bonnet M, Dausset C, Gachinat M, Camarès O, Chassard C, Nivoliez A, Bornes S. 2019. *Lactobacillus rhamnosus* Lcr35 as an effective treatment for preventing *Candida albicans* infection in the invertebrate model *Caenorhabditis elegans*: First mechanistic insights. PLoS One 14.

47. Brenner S. 1974. The genetics of *Caenorhabditis elegans*. Genetics 77:71–94.

48. Mitchell DH, Stiles JW, Santelli J, Rao Sanadi D. 1979. Synchronous growth and aging of Caenorhabditis elegans in the presence of fluorodeoxyuridine. J Gerontol 34:28–36.

49. Wu D, Rea SL, Yashin AI, Johnson TE. 2006. Visualizing hidden heterogeneity in isogenic populations of *C. elegans*. Exp Gerontol 41:261–270.

50. Tsuru A, Hamazaki Y, Tomida S, Ali MS, Komura T, Nishikawa Y, Kage-Nakadai E. 2021. Nonpathogenic *Cutibacterium acnes* confers host resistance against *Staphylococcus aureus*. Microbiol Spectr 9.

51. Liao Y, Smyth GK, Shi W. 2014. featureCounts: an efficient general purpose program for assigning sequence reads to genomic features. Bioinformatics 30:923–930.

52. Anders S, Huber W. 2010. Differential expression analysis for sequence count data. Genome Biol 11.

53. Yu G, Wang LG, Han Y, He QY. 2012. clusterProfiler: an R package for comparing biological themes among gene clusters. OMICS 16:284–287.

54. Pfaffl MW. 2001. A new mathematical model for relative quantification in real-time RT-PCR. Nucleic Acids Res 29:E45.

